# EGFR-targeted and MMP-activated membranolytic peptides derived from *Polybia paulista* MP1 kill cancer cells specifically *in vitro* and reduce tumour growth *in vivo*

**DOI:** 10.1101/2025.10.07.680850

**Authors:** Arindam Pramanik, Andrew Booth, Dagmara Kobza, William J Brackenbury, Simon D Connell, Paul A Beales, Thomas A Hughes

## Abstract

Membranolytic peptides have demonstrated potential as cancer therapeutics, although targeting to cancer cells and reducing toxicity associated with activity in normal cells remain unmet challenges. We have investigated the membranolytic peptide MP1, from *Polybia paulista*, in order to assess ways to reduce its non-specific toxicity and thereby improve its characteristics as a cancer therapeutic. Using a panel of human breast cell lines and cell survival assays, we show that C-terminal addition of an EGFR binding sequence, with or without a linking MMP-2 cleavage sequence, generally reduced the efficacy of the peptides relative to wildtype MP1, as determined by increases in IC_50_ values. Critically, cell lines that show the highest sensitivities to these fusion peptides (MDA-MB-468, MDA-MB-231) expressed the highest EGFR and/or MMP-2 levels, demonstrating that these additions direct the cytotoxic activity to cells expressing these biomarkers. MMP-2 inhibition significantly reduced the cell-killing activity of peptides containing MMP-2 cleavage sites, further demonstrating targeting to this biomarker. Fusion peptides significantly induced apoptosis and reduced survival in EGFR/MMP-2 high cancer cells, while sparing EGFR/MMP-2 low cells in the context of standard tissue culture and 3D-spheroids. Finally, systemic treatment with the EGFR and MMP-2 cleavage fusion significantly reduced tumour size in MDA-MB-468 xenograft models, confirming *in vivo* efficacy against cancer cells and acceptable systemic toxicity. We present this EGFR-MMP-MP1 peptide as a novel cancer therapeutic for further pre-clinical and clinical development.

## Introduction

The majority of cancer therapeutics directly or indirectly target the same cancer property, namely aberrant growth. This is the main mechanism of action for essentially all cytotoxic chemotherapies (1), which inhibit cell cycle processes, and many small molecule inhibitors (2) and biologics (3), which typically target proteins that promote growth. Although these therapies have been successful, newer developments have increasingly focused on alternative mechanisms of action. A potential alternative therapeutic strategy is to lyse the plasma membranes of cancer cells specifically, targeting the differences in lipid composition (4), membrane structure (5), and protein expression (6) that cancer membranes display compared to normal cell membranes. Such lysis would potentially have anti-cancer activity both by directly reducing cancer cell survival, and by inducing immune recognition of cancers through necrotic release of tumour antigens (7). Much of the work in this area has focused on harnessing naturally-occurring membranolytic peptides, sourced from venoms and toxic secretions from a wide range of organisms (8). However, despite development and testing of many different peptides, none have yet entered routine use in the clinic although some clinical trials are on-going (8). Surprisingly, many membranolytic peptides have been reported to exhibit intrinsic cancer-specific activity (9,10), although these reports have not always been confirmed in more extensive studies and this may represent a key limitation for clinical translation as cancer therapies. An example is the peptide MP1 from the wasp *Polybia paulista*; this was initially reported as having some degree of cancer-specific lytic activity (11,12), although our recent work has failed to support this when using a larger panel of cancer and non-cancer cell types (13). Nevertheless, the potential of these highly toxic lytic peptides as oncological therapies remains, if they can be directed specifically to cancer membranes.

In this work, we have targeted the activity of *Polybia paulista* MP1 to cancer cells by adding two different functionalities to its sequence. First, we have taken advantage of a peptide designed to bind specifically to the extracellular domain of the epidermal growth factor receptor (EGFR) (14). EGFR is over-expressed in a range of common cancers, including breast (15,16), colorectal (17,18), and lung (19), and is well established as both a target for therapeutic inhibition (20) and a surface biomarker to direct binding of therapeutics to cancer cells (21-23). Secondly, we have separated the EGFR-binding peptide from MP1 using a sequence that is cleaved by matrix metalloproteinase 2 (MMP-2) (24). MMP-2 is frequently over-expressed in cancers, including in breast and colorectal, and expression is often associated with aggressive features and with poor survival (25-27). Accordingly, MMP-cleavage sites have been used to confer cancer-specific activation properties on various potential therapeutics (24,28,29). We demonstrate that these additions combine to generate a fusion peptide that directs the lytic activity of MP1 to breast cancer cells expressing both targeted biomarkers. The resulting therapeutic peptide has low non-specific toxicity so it can be delivered systemically, and targets cancer cells effectively so as to reduce tumour growth *in vivo*. Thereby, we report a template for further development of a completely novel class of membranolytic cancer therapeutics.

## Materials and methods

### Materials

Peptides were synthesised by Bio-Synthesis Inc (Lewisville, TX, USA). DOPC (1,2-dioleoyl-sn-glycero-3-phosphocholine) was obtained from Avanti Polar Lipids Inc (Alabaster, AL, USA). NaCl and HEPES buffer were obtained from Sigma-Aldrich (St Louis, MA, USA).

### Cell lines

MDA-MB-468, MDA-MB-231, BT-474, AU-565, HB2, and MCF-10A cell lines were procured from the American Type Culture Collection (ATCC) and validated for lack of mycoplasma contamination (MycoAlert; Lonza, Basel, Switzerland) and identity (short tandem repeat profiling; Source Bioscience, Nottingham, UK) prior to experimental use. Cells (except MCF-10A) were cultured in Dulbecco’s Modified Eagle Medium (DMEM) supplemented with 10% (v/v) fetal calf serum (FCS) and 100 units/ml penicillin-streptomycin (all from Thermo Scientific; Waltham, MA, USA). MCF-10A was cultured in DMEM supplemented with 5% horse serum, 0.1μg/ml cholera toxin, 0.5μg/ml hydrocortisone, 0.02μg/ml epidermal growth factor, 100 units/ml penicillin-streptomycin (all from Thermo Scientific, Waltham, MA, USA). Cells were grown at 37°C in a humidified incubator with 5% CO_2_. Cells were maintained and experiments were performed under conditions ensuring cell densities that supported exponential growth.

### Cell survival assays

MDA-MB-468, MDA-MB-231, BT-474, AU-565, HB2, MCF-10A cells were seeded in 96-well plates at 1×10^4^ cells/well in complete growth media and incubated for 18h. Cells were then treated with peptides (0–250µM) for up to 24h. Cells were then incubated with 0.5mg/ml MTT dissolved in PBS for 3h. The formazan crystals formed within the cells were dissolved using 500µl of isopropanol and absorbance was measured at 570nm on a microplate spectrophotometer (Biotech Instruments, USA). For spheroids assays, MDA-MB-468 or HB2 cells (1000 cells/well) were seeded in 250µl of DMEM supplemented with 10% FCS and 2.5% Matrigel (Corning, New York, USA) in low-adherent, round-bottom 96-well plates (Corning, New York, USA). The plates were centrifuged at 360xg for 10min and subsequently incubated for 48h to allow spheroid formation. MDA-MB-468 or HB2 spheroids were then treated with peptides at the IC_50_ doses determined for MDA-MB-468 cells for 24h. Cellular viability within the spheroids was assessed by staining with Hoechst 33342 (5µg/ml) for 30min, followed by propidium iodide (1.5µg/ml) for 10min. Red fluorescence, indicated non-viable cells stained with propidium iodide, whereas blue fluorescence represented both viable and non-viable cells stained with Hoechst 33342. The survival of spheroids was determined by calculating the ratio of blue:red fluorescence using ImageJ software.

### Western blotting

Cells were seeded into 6-well plates at 3×10^5^ and were cultured for 24h. Cells were then harvested by scrapping, washed with ice-cold PBS, and incubated on ice for 45min in RIPA lysis buffer (Thermo Scientific, USA) with agitation to extract whole-cell proteins. Lysates were centrifuged at 13,000rpm for 10min at 4°C, and protein concentrations were determined using a BCA assay (Merck, USA). Proteins were mixed with 2xLaemmli buffer, heated at 90°C for 5min, and resolved by electrophoresis on 4–12% polyacrylamide gels (Bio-Rad, USA). Next, proteins were transferred to PVDF membranes and blocked in 5% skimmed milk in Tris-buffered saline with Tween 20. Membranes were then probed with EGFR or β-actin primary antibody (1:1000; Cell Signalling Technology, USA) overnight at 4°C, and secondary antibodies (1:5000; Cell Signalling Technology, USA) for 2 h. Bands were visualized with Pierce™ ECL reagent (Thermo Scientific, USA), quantified via Chemi-doc (Bio-Rad, USA), and analyzed using ImageJ (NIH, USA).

### Large unilamellar vesicle leakage assays

These assays were performed exactly as previously described (13). In brief, DOPC solution (1ml, 15mM) was dried under a steam of nitrogen, to a thin film, further dried under vacuum, and rehydrated in 5(6)-Carboxyfluorescein (CF) solution (120mM in 10mM HEPES pH 7.4) followed by 5 freeze-thaw cycles in liquid nitrogen. The suspension was then extruded through a 400nm polycarbonate membrane and unencapsulated CF was removed by size exclusion chromatography. Leakage assays were carried out using a Hamilton Microlab Star M liquid handling robot (Hamilton robotics Ltd.) using a serial dilution of peptide into 10mM HEPES (pH 7.4) buffer, followed by a fixed concentration of CF-loaded vesicles. Negative and positive controls were established by addition of vesicles to peptide-free buffer and buffer containing 0.6mM Triton X-100. The final assay plate (384-well black OptiPlate, PerkinElmer LAS (UK) Ltd) was transferred to a Perkin Elmer Envision plate reader where the CF fluorescence intensity was measured (ex: 495 nm /em: 517 nm).

### MMP-2 analyses

Matrix metalloproteinase-2 (MMP-2) was analysed in all 6 cell lines. Cells were seeded in 6-well plates at 3×10^5^ in complete medium and allowed to grow for 48h. 500μl of cell media from each cell line was collected and centrifuged at 5000rpm for 15min at 4°C to remove any cells or debris. The supernatant was collected and 50μl of each culture media was added in 3 replicate wells of a monoclonal mouse anti-human MMP-2 antibody pre-coated 96-well ELISA plate (Human Total MMP-2 kit; BioLegend, USA). It was then mixed with the respective assay buffers as per the kit instruction. The plate was then sealed and incubated at room temperature for 2h. The supernatant sample was then discarded and the wells were washed with assay buffer several times. Next, 100μl of Human Total MMP-2 Detection Antibody solution was added into the wells and incubated for 1h. 100μl Avidin-HRP solution was then added for 1h and finally, after washing, 100μl substrate solution was added.

Absorbance was measured at 450nm (Biotech Instruments, USA). The final MMP-2 concentration was calculated against a standard curve. For inhibitor studies, MDA-MB-468 cells were seeded in 96-well plates at 1×10^4^ cells/well in complete growth media and incubated for 18h. Cells where then treated with MMP-2 inhibitor (2μM chlorhexidine dihydrochloride; Santa Cruz Biotechnology, USA), peptide (20μM EGFR-MP1 or 30μM EGFR-MMP-MP1), or the combination for 24h. Post-treatment, MTT assays were performed as detailed above.

### Apoptosis analysis using flow cytometry

Apoptosis was studied using Annexin V/propidium iodide (PI) assays and flow cytometry. In brief, MDA-MB-468 and HB2 cells were seeded in 6-well plates at 3×10^5^ in complete medium and allowed to grow for 18h. Cells were then treated with the IC_50_ doses for MDA-MB-468 cells of EGFR-MP1, EGFR-MMP-MP1 or EGFR-MMP-D2K for 24h. Following treatment, the cells were scraped from the plate and washed with Annexin binding buffer. Then 2µg/ml Annexin V-FITC (Thermo Fisher Scientific, Waltham, MA, USA) was added and incubated for 15min in the dark. Cells were then washed and 1µg/mL propidium iodide (PI, Thermo Fisher Scientific, Waltham, MA, USA) was added. Then cells were analyzed using a CytoFLEX S flow cytometer (Beckman Coulter, UK), and the data were processed using FlowJo software (v10.6.1).

### Ex vivo *haemolysis assay*

Blood was collected from five 10-week-old male C57BL/6 mice via cardiac puncture following cervical dislocation, using paediatric blood collection tubes containing K3 EDTA (Greiner; Slušovice, Czech Republic). The whole blood was then centrifuged at 500xg for 5min at 4°C, and the plasma was carefully removed. The remaining red blood cells (RBC) were gently resuspended to their original volume using 150mM NaCl. RBC were repeatedly washed and diluted to 1:50 in PBS. For a positive control, 1% (v/v) Triton X-100 was used (100% lysis), while cells incubated in PBS served as a negative control (0% lysis). 200μL of RBC in triplicate was treated with concentrations (25, 50 or 100μM) of MP1, EGFR-MP1, EGFR-MMP-MP1, EGFR-D2K or EGFR-MMP-D2K and incubated on an orbital shaker at 100rpm for 1h at 37°C. Following incubation, the samples were centrifuged at 500xg for 5min to collect the intact cells. The supernatants, containing lysed RBCs, were collected and hemolysis was quantified by measuring absorbance at 540nm.

### In vivo *studies*

These were performed by HD Biosciences (Shanghai, China). All procedures related to handling, care, and treatment of animals were performed according to guidelines approved by the Institutional Animal Care and Use Committee (IACUC, #AUC105) of HD Biosciences, under the criteria of the Association for Assessment and Accreditation of Laboratory Animal Care (AAALAC). Protocols and AAALAC accreditation were reviewed by the Animal Welfare and Ethical Review Committee at the University of Leeds, and were approved (reference THAWERC222707; date 28/7/2022). Protocols and data are reported in accordance with the ARRIVE guidelines (https://arriveguidelines.org/arrive-guidelines). Female NCG mice (6-8 weeks old) were purchased from Gempharmatech Ltd (Nanjing, China). Triple negative breast cancer xenografts were established by subcutaneously injecting exponentially growing MDA-MB-468 cells (1×10^7^ cells in 0.2ml in D-PBS 1:1 with Matrigel) into the right flank of the mice. Once the tumour volumes reached 150–200 mm^3^, mice bearing xenografts were assigned into two groups using stratified randomization based upon tumor volumes in Microsoft Excel to ensure groups were comparable at baseline. 2 groups were used: control group (13 animals) and the test group (7 animals); group sizes were not determined by power calculation due to lack of data on which this could be based. The test group was administered three intravenous doses via tail vein injection of 500µg EGFR-MMP-MP1 on days 1, 3, and 5, while the control group received equivalent volumes of saline. Tumour volumes were monitored using callipers, and body weight was monitored throughout the duration of experiment; no animals were excluded. Experimenters were not blinded to treatment group. The experiment was terminated on day 27 and the tumour tissue was extracted and weighed.

### Statistics, on-line data, and data availability statement

*In vitro* experiments were performed as biological triplicates unless stated otherwise in figure legends. Statistical analyses, as described in figure legends, were performed using Prism v10 (Graphpad; Boston, MA, USA). The Depmap portal data (Fig S4A) were accessed at https://depmap.org/portal/gene/EGFR?tab=dependency&characterization=expression as transcripts per million averaged across replicates of each sample. The Protein Atlas portal data (Fig S4B) were accessed at https://www.proteinatlas.org/ENSG00000146648-EGFR/cell+line#breast_cancer as normalized transcripts per million. Essentially all data are included within the figures, although primary data can be requested from the authors without restriction.

## Results

### MP1 toxicity and cell-line specificity can be modified using C-terminal extensions

We previously reported that MP1 is a membranolytic peptide capable of causing cell death in human cells with IC_50_ doses varying across a panel of cell lines by over 6-fold without specificity for cancer cells (13) (Fig S1). Our first aim was to assess whether MP1 could gain cancer specificity by extending the peptide sequence with additions known to induce binding to the surface of cancer cells, or to allow activation by cancer cells. We investigated addition of a targeting sequence that directs binding to EGFR (14). Also, we linked this to MP1 either by a simple GG linker or using a proteolytic cleavage site that can be targeted by MMP-2 (24). We attached these targeting or cleavage sequences to either the N-terminus or the C-terminus of MP1. See Table 1 for the sequences used and our nomenclature for the different peptides. A panel of human breast epithelial cell lines was established, including two lines from a non-transformed origin (MCF-10A; HB2) and four cancer lines (BT-474; AU-565; MDA-MB-231; MDA-MB-468). Cells were treated with different doses of MP1 or the modified peptides. Survival was assessed relative to untreated using MTT assays (Figs S2 and 1) and IC_50_ doses were determined for each peptide in each cell line.

**Table 1.**
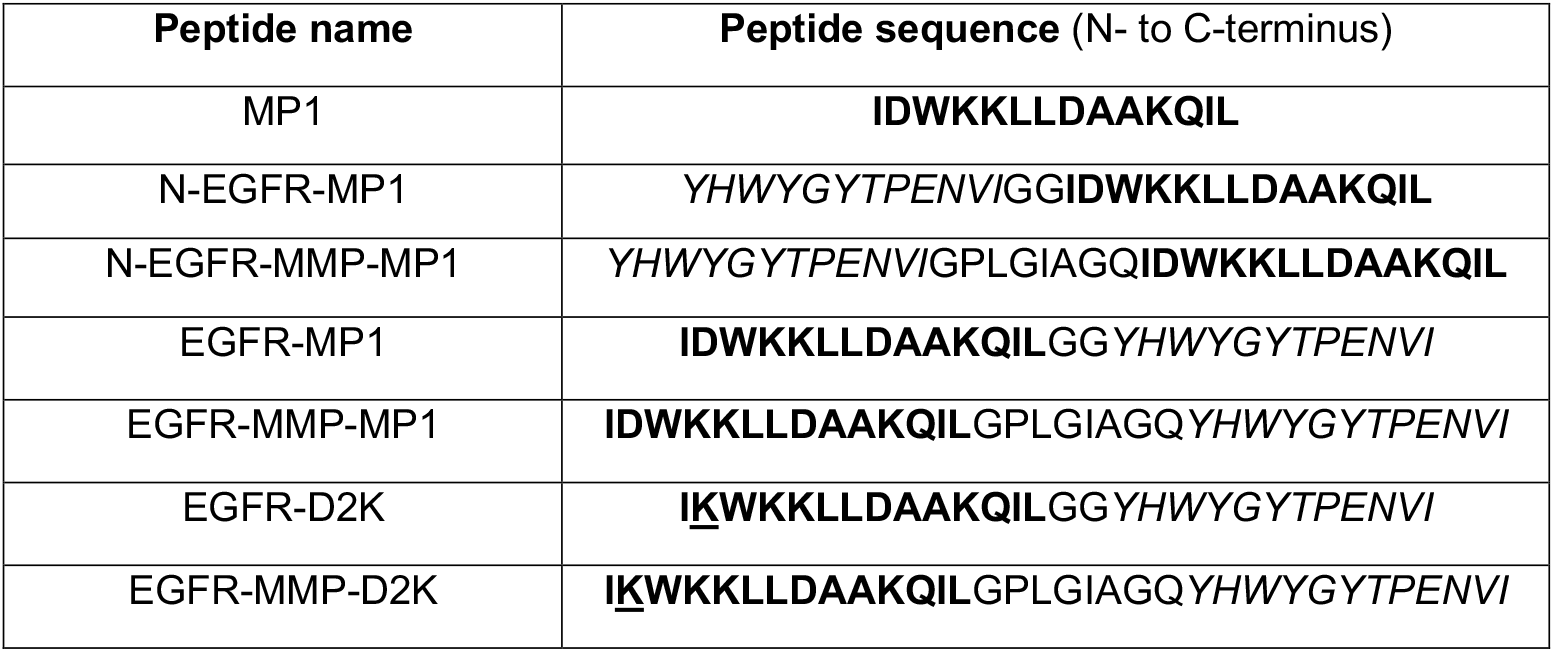
Peptide sequence details. The MP1 sequence is in bold; substitutions are underlined; the EGFR-targeting sequence is in italics.

We found that N-terminal fusions had very limited activity, with IC_50_ doses estimated to be up to 10-times higher when compared to MP1 itself (Fig S2). We concluded that an unmodified N-terminus was required for MP1 function, and that these fusions had little potential as anti-cancer agents; these were not investigated further. By contrast, the C-terminal fusions showed a range of effects that varied according to cell line (Fig 1). In most cell lines EGFR-targeting reduced efficacy, although the extent of this reduction varied from negligible (MDA-MB-231) to 5-fold (AU-565). Similarly, the addition of the MMP cleavage site further reduced efficacy. MCF-10A cells, however, behaved differently; in this cell line neither fusion peptide showed strikingly different efficacy when compared to MP1 (Fig 1A), although it should be noted that this cell line is especially resistant to the wildtype MP1 peptide and therefore further reductions in function in an already poorly-functional peptide may be difficult to detect. We concluded that the C-terminal fusion peptides demonstrated activities that were dependent on the characteristics of the cells used, and therefore that their dependence on EGFR expression and MMP activity should be investigated.

**Figure 1.**
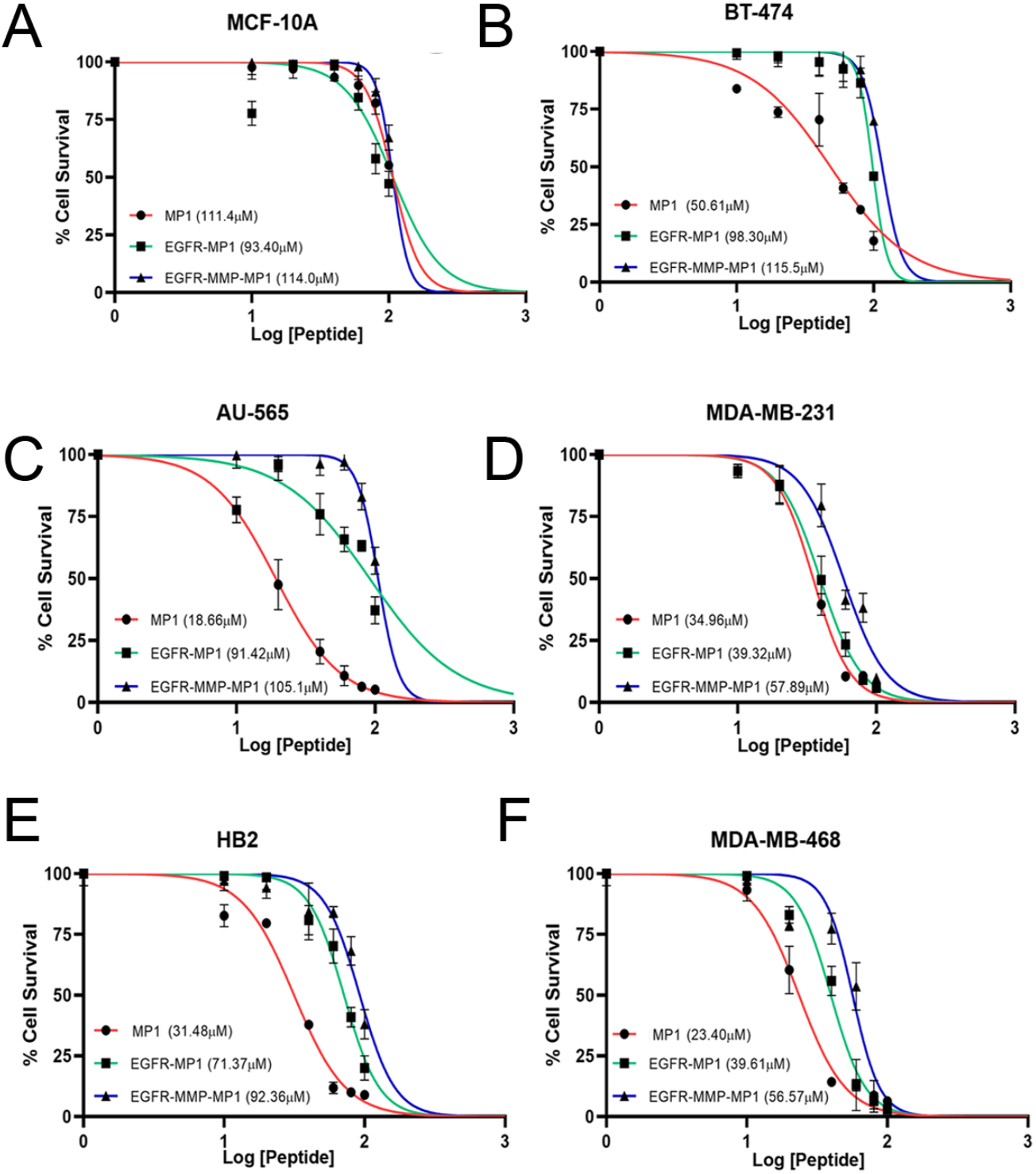
Addition of EGFR-targeting and/or MMP-2 cleavage sequences to MP1 modifies lytic activity in a cell-specific manner. Cell lines as marked were treated with a range of doses of MP1, EGFR-MP1 or EGFR-MMP-MP1 for 24h and cell survival was assessed using MTT assays. Survival is shown relative to untreated control and IC_50_ values were extracted from best-fit curves. Data represent means and standard errors of three independent experiments.

We also assessed the activity of MP1 and EGFR-MMP-MP1 against simple model membrane vesicles, to demonstrate that the peptides act directly on the membrane in a system where down-stream biology cannot be induced. Vesicles were assembled using the lipid DOPC and were loaded with the carboxyfluorescein at concentrations that are self-quenching with respect to fluorescence. Vesicles were treated with a range of concentrations of peptides, and fluorescence resulting from release of carboxyfluorescein and consequent loss of quenching was quantified (Fig S3). As expected, MP1 was highly effective at lysing vesicles, which can be quantified using the concentration required to achieve 50% of maximal fluorescence: nM. In comparison, EGFR-MMP-MP1 showed a very large reduction in efficacy (more than 18-fold; 50% lysis concentration 1.39µM), which most likely reflects the complete lack of EGFR and MMP-2 in this purified system.

### Activity of the targeted and cleavable peptides is dependent on expression of EGFR and MMP-2

To correlate the cytotoxic efficacy of fusion peptides with EGFR expression, we quantified EGFR in the panel of cell lines using Western blots (Fig 2A). The cell lines divided broadly into two groups: MDA-MB-468 and MDA-MB-231 cells had similar, relatively high, expression levels while the remaining lines had low levels. RNA expression data for the cancer lines, available from two independent resources (DepMap Portal and Human Protein Atlas), also confirmed the highest EGFR expression in MDA-MB-468 cells, followed by MDA-MB-231 cells (Fig S4). In accordance with their high EGFR expression, these two lines showed the greatest sensitivity to EGFR-MP1 across the cell line panel (Fig 1), and also showed comparatively small reductions in efficacy associated with the addition of EGFR-targeting when compared to wildtype MP1 itself. This is in contrast to, for example, AU-565 that had the greatest intrinsic sensitivity to wildtype MP1 but showed striking and substantial reductions in efficacy from EGFR-targeting in accordance with its low EGFR expression.

**Figure 2.**
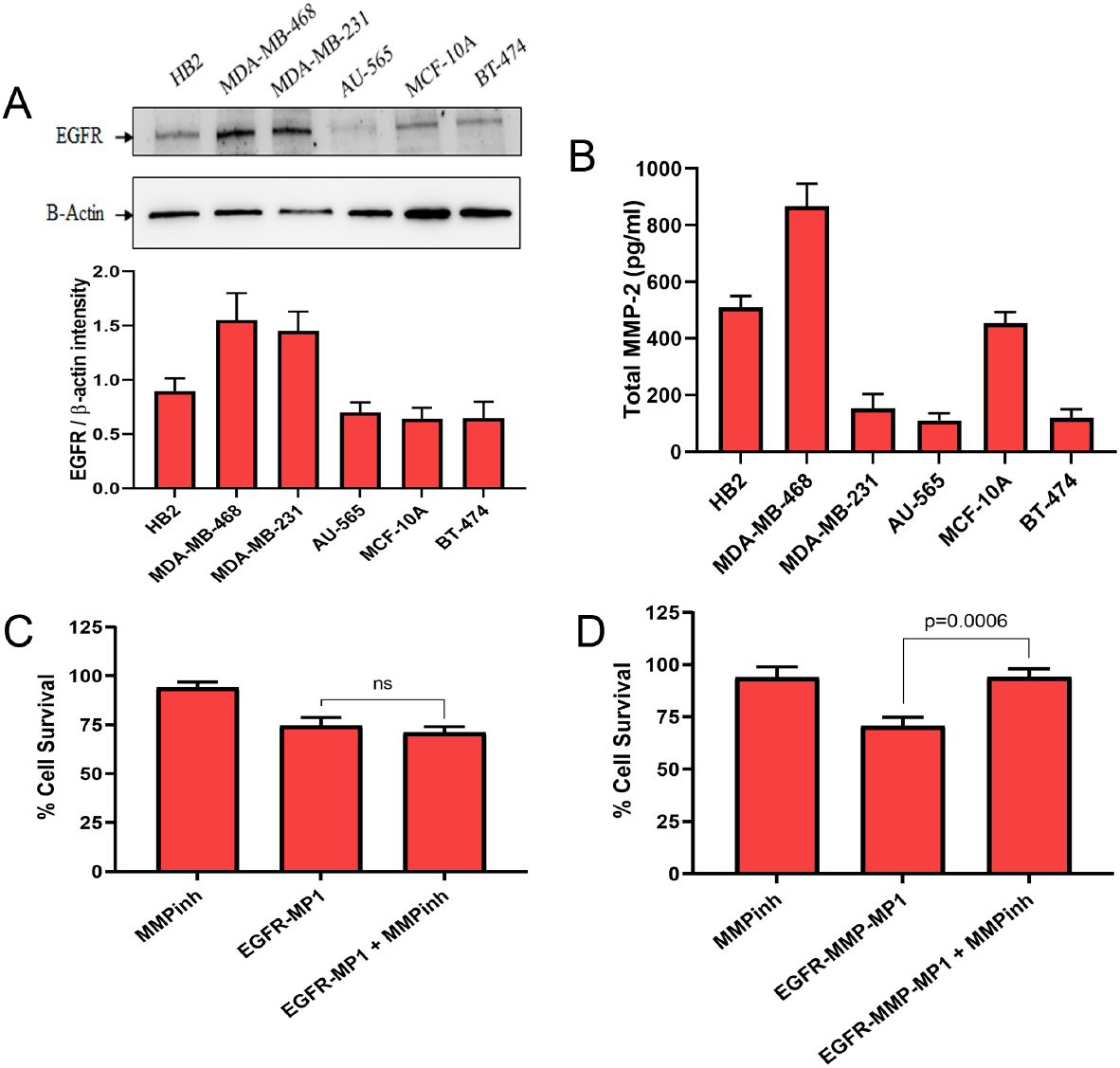
MDA-MB-468 cells express high levels of EGFR and MMP-2, and show MMP-2 dependent activity of EGFR-MMP-MP1. (A) EGFR expression in breast epithelial cell lines was assessed using western blots (top) and expression was quantified relative to actin using densitometry (bottom). (B) Soluble MMP-2 was quantified in the medium of cultured cells by ELISA. (C and D) MDA-MB-468 cells were treated with 20μM EGFR-MP1 (C) or 30μM EGFR-MMP-MP1 (D) in the presence or absence of 2μM of the MMP-2 inhibitor chlorhexidine dihydrochloride (MMPinh), or with the inhibitor alone. Cell survival was measured using MTT assays and is shown relative to an untreated control. Quantitative data represent means and standard errors of three independent experiments. Statistical analyses were performed using a one-way Student’s T test.

Next, we quantified expression of MMP-2 in the panel of cell lines using ELISAs (Fig 2B). MDA-MB-468 demonstrated the highest secreted concentration, which was more than 4-fold higher than the remaining cancer lines, with the two non-transformed lines (HB2; MCF-10A) showing intermediate activities. Accordingly, MDA-MB-468 cells showed relatively high sensitivity to the targeted and cleavable peptide EGFR-MMP-MP1, as – surprisingly – did MDA-MB-231 cells (Fig 1) despite relatively low MMP-2 activity. AU-565 cells had the lowest MMP-2 expression (Fig 2B) and also showed the greatest loss of efficacy associated with the addition of the MMP cleavage site, indicating that MP1’s efficacy was suppressed in this fusion when MMP-2 levels were low.

We were also interested to test more formally whether the efficacy of EGFR-MMP-MP1 was dependent on MMP-2 activity as would be expected. Therefore, we treated MDA-MB-468 cells with EGFR-MMP-MP1 or with EGFR-MP1, in the presence or absence of the MMP-2 inhibitor chlorhexidine dihydrochloride, and assessed cell survival relative to untreated (Figs 2C and 2D). Inhibition of MMP-2 activity alone caused a small and non-significant decrease in cell survival, while -as expected – both peptides reduced cell survival significantly. Inhibition of MMP-2 activity completely halted the cell death induced by EGFR-MMP-MP1 (Fig 2D; p<0.001) while it had no effect on the activity of EGFR-MP1 (Fig 2C), thereby demonstrating that the MMP-2 cleavage site confers MMP-dependent activity on the peptide.

We concluded that the efficacy of the fusion peptides correlated with expression or activity of the markers they were designed to target, with MDA-MB-468 cells showing particularly favourable characteristics for successful targeting by this combination.

### Inclusion of EGFR-targeting and MMP-cleavage confers a therapeutic window to target EGFR and MMP-2 positive cells

Next, we aimed to assess whether the EGFR-targeting and MMP-dependence could give sufficient specificity to our peptides to kill target cells (cancer cells that are EGFR and MMP-2 positive) while sparing non-target cells (cells with low expression of both or either marker). This analysis is potentially confounded by differences in intrinsic sensitivity to wildtype MP1, highlighted for example by the relative resistance to all MP1-derived peptides seen in MCF-10A and BT-474 cells that was unrelated to EGFR or MMP expression. We selected MDA-MB-468 cells as our ideal target cell since it had the highest expression of both EGFR and MMP-2 (Fig 2). For comparison, we selected HB2 cells as a representative non-target cell since these cells uniquely had intrinsic sensitivity to wildtype MP1 that was similar to MDA-MB-468 cells (HB2 IC_50_ 31µM, compared to MDA-MB-468 23µM; Fig 1), but also had lower expression of both targeting markers (see Fig 2).

HB2 or MDA-MB-468 cells, in either standard 2D culture or cultured as 3D spheroids, were treated with the IC_50_ doses of either EGFR-MP1 or EGFR-MMP-MP1 as determined in Fig 1 for MDA-MB-468 cells, or were treated with control. Cell death in these cultures was assessed in the 2D cultures by annexin-V/propidium iodide staining and flow cytometry (Fig 3A; Fig S5 for representative cytometry plots). Cell survival in spheroids was assessed by Hoechst 33342/propidium iodide staining and microscopy (Fig 3B; Fig S6 for representative microscopy images). We found that both peptides induced substantial apoptosis and reduced cell survival in MDA-MB-468 cells, while effects on HB2 cells were negligible. We concluded that our EGFR-targeting and MMP-activation strategy was sufficient to allow specific killing of the target cells.

**Figure 3.**
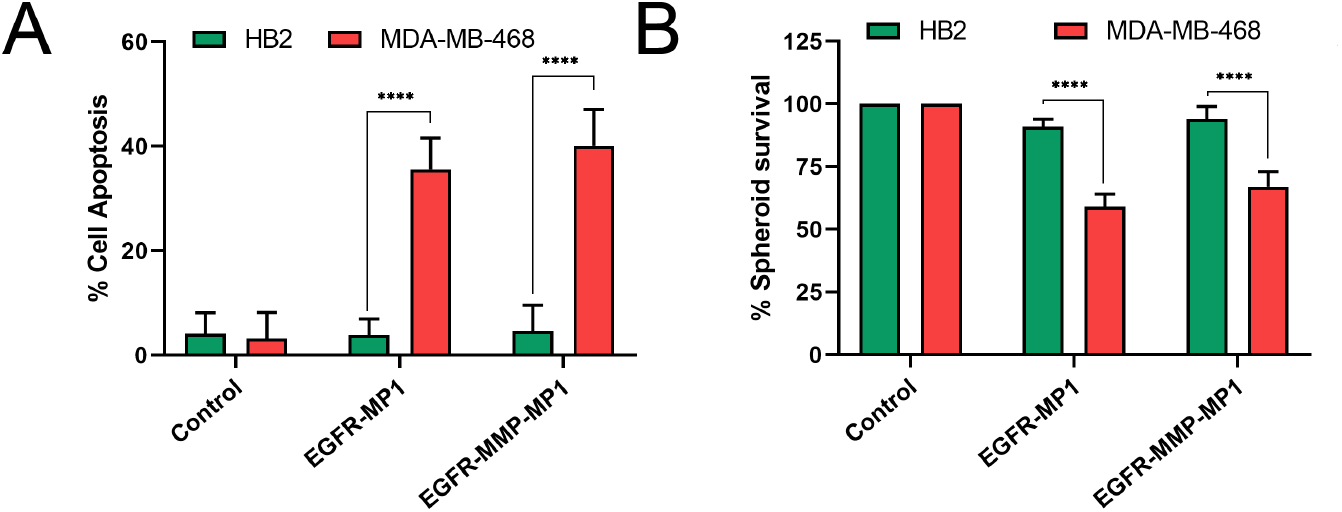
Directing MP1 to EGFR and MMP-2 allows targeting of MDA-MB-468 cancer cells, sparing non-cancerous HB2 cells. (A) HB2 or MDA-MB-468 cells in 2D culture were treated with EGFR-MP1 (39.6μM) or EGFR-MMP-MP1 (56.6μM) for 24h. Apoptosis was quantified using Annexin V/PI staining and flow-cytometry. (B) 3D spheroids were established with HB2 or MDA-MB-468 cells, and these were treated with EGFR-MP1 (39.6μM) or EGFR-MMP-MP1 (56.6μM) for 24h. Cell survival was quantified by counting fluorescent cells after staining with PI/Hoechst 33342 by fluorescence microscopy. Data represents means and standard errors of three independent experiments, and statistical analyses were performed using two-way ANOVA tests (**** indicates p<0.0001).

### Substitutions within the MP1 sequence can increase toxicity

We have previously reported on changes in MP1 activity caused by substitution of individual residues within its sequence (13). We were now interested to assess whether any of these substitutions could improve the efficacy and/or specificity of our fusion peptides. In Fig 4A, we present a re-analysis of our previous data (13) demonstrating changes in IC_50_ values associated with four separate single residue substitutions. We show that in MDA-MB-468 cells, increased efficacy in terms of cell killing (ie reduced IC_50_ values) results from substitutions where the aspartic acid residues at position 2 or 8 are replaced with lysines (D2K and D8K respectively). However, we noted that D8K also showed increased efficacy in MCF-10A cells, which for our current purposes represents a ‘non-target’ cell line. Therefore, we selected D2K as the substitution with most potential to improve efficacy and specificity.

**Figure 4.**
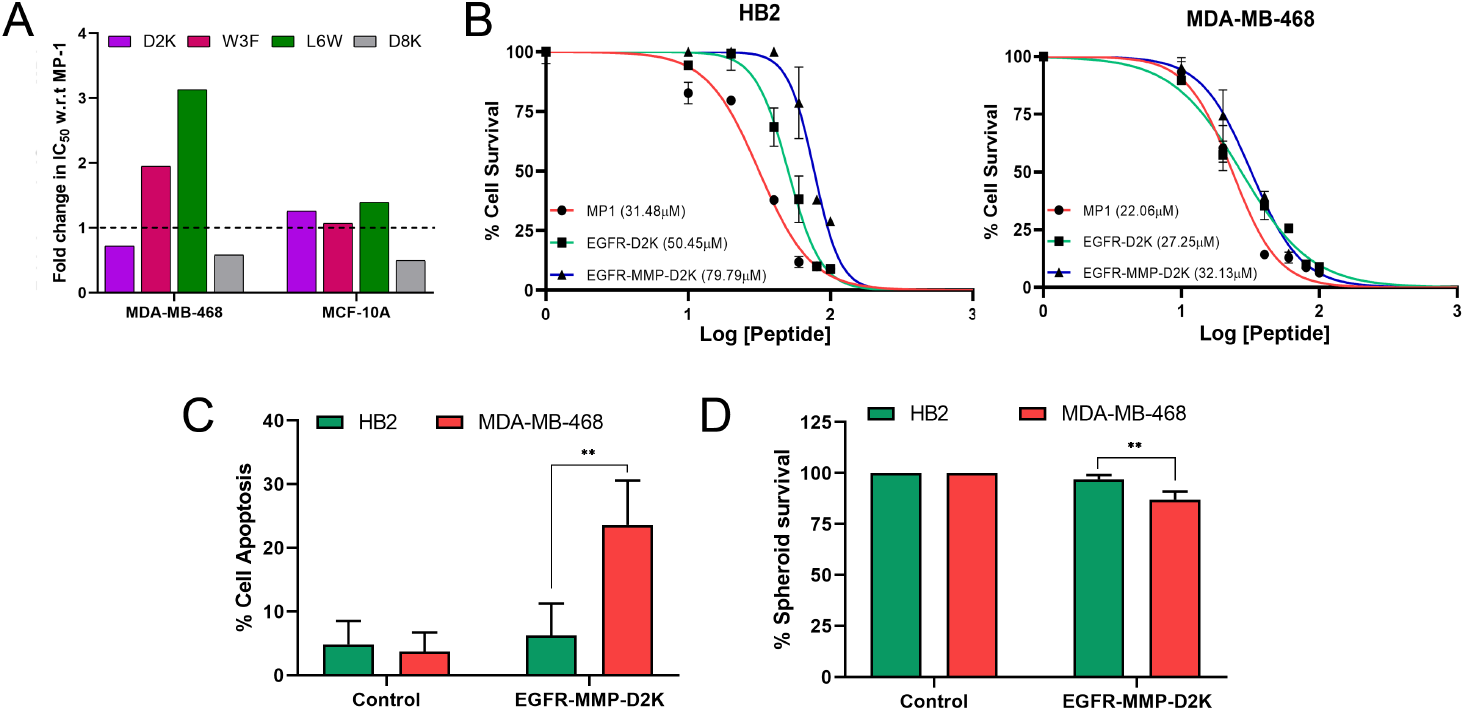
The D2K substitution in MP1 increases toxicity in the context of EGFR-MP1 and EGFR-MMP-MP1, and specificity for MDA-MB-468 cells is retained. (A) IC_50_ values for MP1 variants with single residue substitutions (D2K, W3F, L6W or D8K) were determined in MDA-MB-468 and MCF-10A cells using MTT assays (Booth et al 2025). Data represent fold change in IC_50_ relative to MP1; variants below the dotted line have reduced IC_50_ values and are therefore have improved efficacy. (B) The D2K substitution was synthesised in the context of the fusion peptides, to create EGFR-D2K and EGFR-MMP-D2K. HB2 or MDA-MB-468 were treated for 24h with various doses of EGFR-D2K or EGFR-MMP-D2K, or with MP1 for comparison, and cell survival was quantified using MTT assays relative to untreated. IC_50_ values were extracted from best-fit curves. (C and D) HB2 and MDA-MB-468 cells in 2D (C) or 3D spheroid (D) culture were treated with EGFR-MMP-D2K (32.1μM) for 24h and apoptosis (C) or cell survival (D) was analysed as in Figure 3. Data represents means and standard errors of three independent experiments, and statistical analyses were performed using two-way ANOVA tests (** indicates p<0.01).

The D2K substitution was synthesized in the context of our EGFR- and EGFR-MMP-extended peptides to create EGFR-D2K and EGFR-MMP-D2K (see Table 1). HB2 or MDA-MB-468 cells, our established pair of non-target and target cells, were treated with different doses of wildtype MP1 or with the targeted D2K peptides, and survival was assessed relative to untreated as previously (Fig 4B). In both cell lines, the D2K peptides were substantially more toxic than their MP1 versions (compare to Fig 1E and F), with a mean reduction in IC_50_ of 17.5µM. Disappointingly, there was no suggestion that increased toxicity was greater in the target cell line MDA-MB-468 as opposed to the non-target HB2. Nevertheless, we repeated our assessment as in Fig 3 of whether the targeted peptide provided sufficient specificity to kill MDA-MB-468 cells while sparing HB2 cells (Fig 4C and 4D; Fig S7 and S8 for representative primary data). We found that EGFR-MMP-D2K significantly reduced cell survival in MDA-MB-468 cells while effects on HB2 cells were minimal, demonstrating a therapeutic window for this substituted peptide with increased efficacy.

### EGFR-MMP-MP1 retards tumour growth in vivo

Our next aim was to assess whether our various peptides had potential as anti-cancer therapeutics using an *in vivo* model. To our knowledge, MP1 or MP1-derived peptides have not previously been used experimentally *in vivo*, therefore we first performed an *in vitro* hemolysis assay to aid selection of peptides that could be suitable for intravenous delivery. Red blood cells were isolated from mouse blood and treated with three different doses of MP1 wildtype or our targeted peptides (EGFR and EGFR-MMP versions of both MP1 and D2K) and hemolysis was measured (Fig 5A). Wildtype MP1 caused unacceptably high levels of hemolysis at all doses, highlighting its relative lack of specificity in terms of cell types lysed. All the fusion peptides showed dramatically lower levels of hemolysis, in accordance with their expected targeting to cancer biomarkers. However, D2K variants showed higher levels of hemolysis than their matched MP1 versions at the two lowest doses used, rising to as high as 5.9% at 50µM (EGFR-MMP-D2K). From the available data, we selected EGFR-MMP-MP1 for further investigation *in vivo*, as it demonstrated low hemolysis at relevant doses (Fig 5A), and showed strong efficacy against our target cell line MDA-MB-468 (Fig 3).

**Figure 5.**
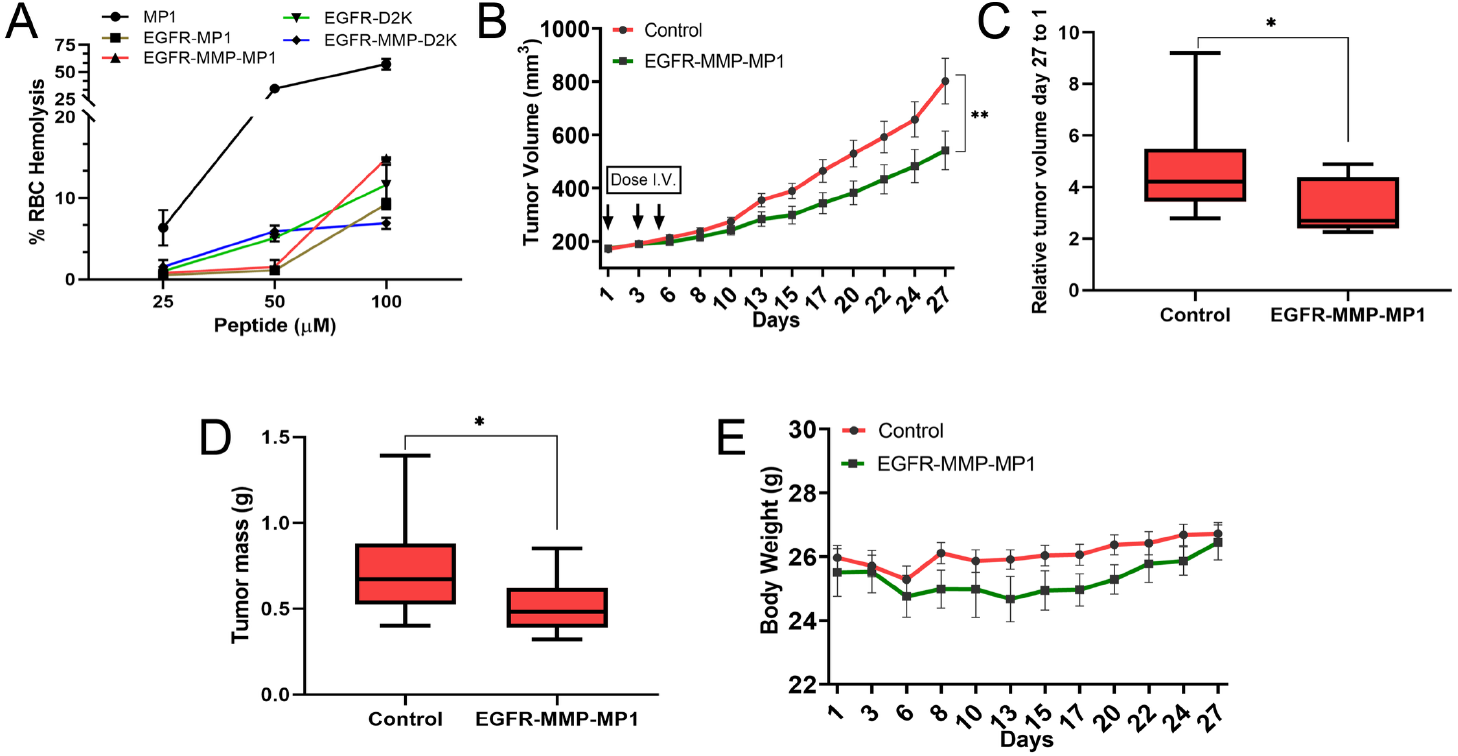
EGFR-MMP-MP1 shows effective anti-cancer activity on MDA-MB-468 xenografts *in vivo*. (A) Hemolysis assays were performed for 5 peptides sequences (MP1, EGFR-MP1, EGFR-MMP-MP1, EGFR-D2K, EGFR-MMP-D2K) in fresh red blood cells isolated from mice. (B-E) MDA-MB-468 xenografts were established in NCG mice before treatment with three doses of 500μg of EGFR-MMP-MP1 peptide or saline control as shown. Tumour size (C) and animal weight (E) was measured at 2-3 day intervals for a total of 27 days. Increase in tumour size over the course of the experiment was quantified as fold-change in size in each group (C). Tumour masses were measured after termination of the experiment (D). Data represent means with standard errors. Statistical analysis was performed using Wilcoxon matched pair signed ranked tests (* indicates p<0.05; ** indicates p<0.01).

NCG immune-compromised female mice were implanted sub-cutaneously with MDA-MB-468 cells. When tumours reached between 150-200mm^3^, animals were randomized to control and treatment groups, and were treated by tail vein injection with doses of control (saline) or 500µg EGFR-MMP-MP1 on days 1, 3 and 5 of the experiment. Tumour size and animal weight were monitored every 2 to 3 days for a total of 27 days, after which animals were sacrificed and tumours were dissected and weighed. Treatment with EGFR-MMP-MP1 caused a significant and sustained retardation in tumour growth (Fig 5B-D). At the end of the experiment, the mean tumour volume of tumours treated with EGFR-MMP-MP1 was only 67.5% (p<0.01) of the untreated tumours, and similarly, tumour mass was only 69.35% (p<0.05). Overall tumour growth from day 1 to day 27 was a 3.1-fold increase in the EGFR-MMP-MP1 treated group, compared to 4.6-fold for the control group (Fig 5C; p<0.05). Animal weights, as a surrogate for side-effects, were slightly reduced in the treatment group, although they recovered by the end of the experiment (Fig 5E). We concluded that EGFR-MMP-MP1 is effective at killing target cancer cells, and is associated with acceptable non-specific toxicity. This novel agent has potential as an anti-cancer therapeutic for EGFR-positive and MMP-2-positive cancers.

## Discussion

Membranolytic peptides have been studied extensively *in vitro* as potential anti-cancer therapeutics (8-10), with a limited number going on to *in vivo* assessment (30,31) or even early-phase clinical trials (32-34). However, toxicity associated with activity against non-cancer cells remains a substantial and mainly unaddressed problem. Investigators have attempted to limit these off-target effects to some extent through delivery by intra-tumoural injection (30-32,34) rather than systemically. Despite this mitigation, agents have still shown substantial toxicity in mouse models (30). Nevertheless, the peptide LTX-315 entered clinical trials as a first-in-class untargeted membranolytic peptide delivered through intra-tumoural injection (32), and has shown some evidence of anti-tumour activity and a toxicity profile that could be tolerable. Further trials of this, and at least two different membranolytic peptides have recently recruited or are underway (33-35). However, use of intra-tumoural injections presents challenges for integration into treatment of many cancers. This is because relatively few cancers are located to allow intra-tumoural injection, and also because the predominant curative regimen for many primary solid cancers prioritises resection surgery, with further therapies usually in the adjuvant (post-surgical) setting. In any event, the main target of systemic therapies for primary solid cancers is often the sub-clinical disseminated cancer cells in unknown locations, which if left untreated can develop into distant metastatic recurrences; intra-tumoral injection is unable to substitute for this role. Consequently, systemic delivery of membranolytic peptides would maximise their potential utility. Critically, however, to allow systemic delivery the issue of specificity to cancer cells needs to be addressed.

We have examined the membranolytic peptide MP1 from the wasp species *Polybia paulista*, which was reported to show intrinsic specificity to cancer cells (11,12). However, we failed to confirm this specificity, demonstrating its activity on human cells to be irrespective of cancer or non-cancer origin (Fig S1). We also found it to have unacceptably high haemolytic activity (Fig 5A), ruling out systemic delivery. Therefore, we focused on extending the MP1 sequence to target it more effectively to cancer cells and spare non-target cells. This approach is related to that taken with the peptide EP-100 (36), which comprises an 18-residue lytic peptide linked directly to a 10-residue sequence that directs binding to a cancer biomarker, the GnRH receptor. This peptide showed lytic activity *in vitro* that was target-specific, and accordingly was safely delivered systemically in mouse models (37). Human clinical trials are on-going, using systemic delivery by intravenous infusion, and initial data show a good safety profile and some anti-cancer activity (38). In our work, we extended the MP1 lytic peptide to include sequences known to bind to EGFR, a very commonly expressed cancer biomarker (20), and to direct cleavage from this targeting sequence by MMP-2, a protease commonly upregulated in cancer cells (29). This strategy was designed to use biomarkers that are over-expressed across a wide range of cancer types including the three commonest solid cancers (for example more widely than the GnRH receptor (39)) and to increase cancer-specificity further by having both targeting and proteolytic activation elements. We have focused on breast cancer, and we found that cell lines from the triple negative sub-classification (MDA-MB-468 and MDA-MB-231) are the most targetable using this biomarker combination (Fig 1 and 2). This is ideal since triple negative primary breast cancer currently lacks molecularly-targeted therapies, has the poorest outcomes, and is most in need of alternative approaches (40). Our data demonstrate that the peptide EGFR-MMP-MP1 is well-tolerated in systemic treatment and causes significant retardation of tumour growth *in vivo* (Fig 5). We believe this novel peptide has great potential for further pre-clinical and clinical development as a cancer therapeutic.

## Acknowledgments

This research was supported by the UK Engineering and Physical Sciences Research Council (EPSRC): EP/R03608X/1.

