## Supplementary material for "EGFR-targeted and MMP-activated membranolytic peptides derived from *Polybia paulista* MP1 kill cancer cells specifically *in vitro* and reduce tumour growth *in vivo*": Figures S1 - S8

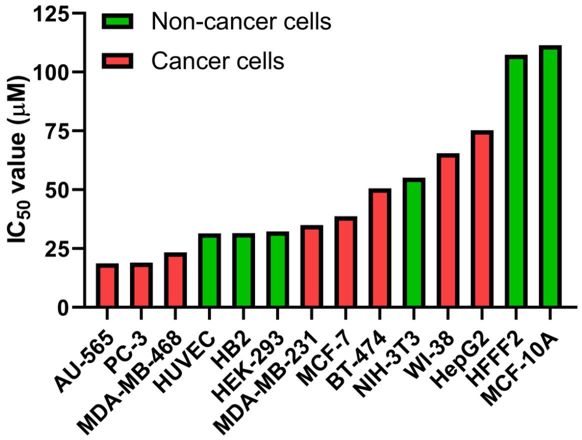


**Figure S1.** MP1 does not show specificity to cancer cells. The $IC_{50}$ doses for treatment with MP1 for 24h were determined in 14 different human cell types using MTT assay; primary data are available in Booth et al 2025. Green bars represent non-cancerous cells while red indicates cancer cells.

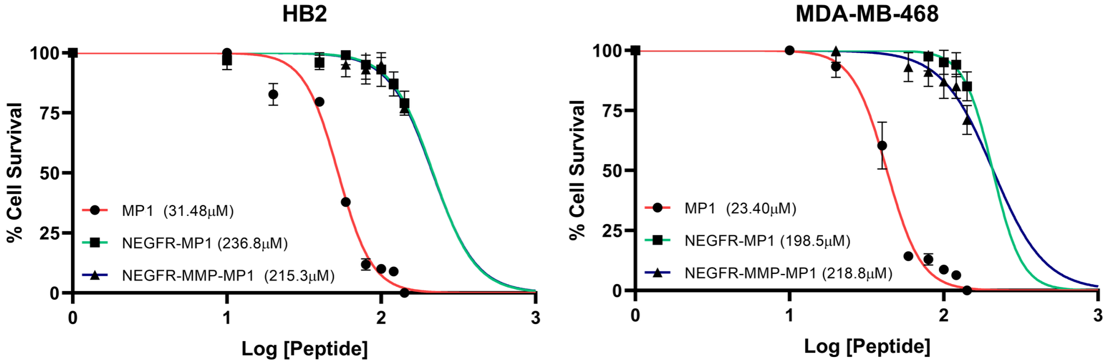


**Figure S2.** N-terminal addition of EGFR- or MMP-targeting to MP1 dramatically inhibits MP1 function. Cell lines as marked were treated with a range of doses of MP1, N-EGFR-MP1 or N-EGFR-MMP-MP1 for 24h and cell survival was assessed using MTT assays. Survival is shown relative to untreated control and $IC_{50}$ values were extracted from best-fit curves. Data represent means and standard errors of three independent experiments.

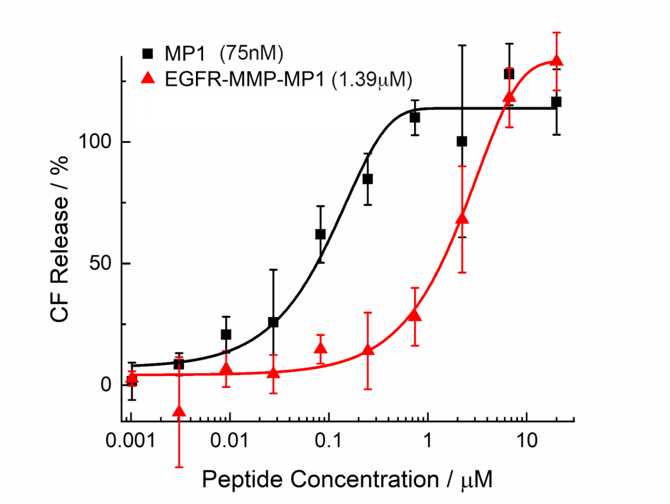


**Figure S3.** MP1 and EGFR-MMP-MP1 act directly on membranes to cause rupture. DOPC large unilamellar vesicles containing carboxyfluorescein at self-quenching concentrations were assembled. These were treated with a range of concentrations of MP1 or EGFR-MMP-MP1 and leakage was quantified by detection of fluorescence associated with release of carboxyfluorescein from the vesicles. Data represent means of replicate wells with standard errors, and are shown as a percentage of complete carboxyfluorescein release as defined by treatment with the detergent Triton-X100. Doses of MP1 or EGFR-MMP-MP1 required to achieve 50% maximal fluorescence are shown in the legend.

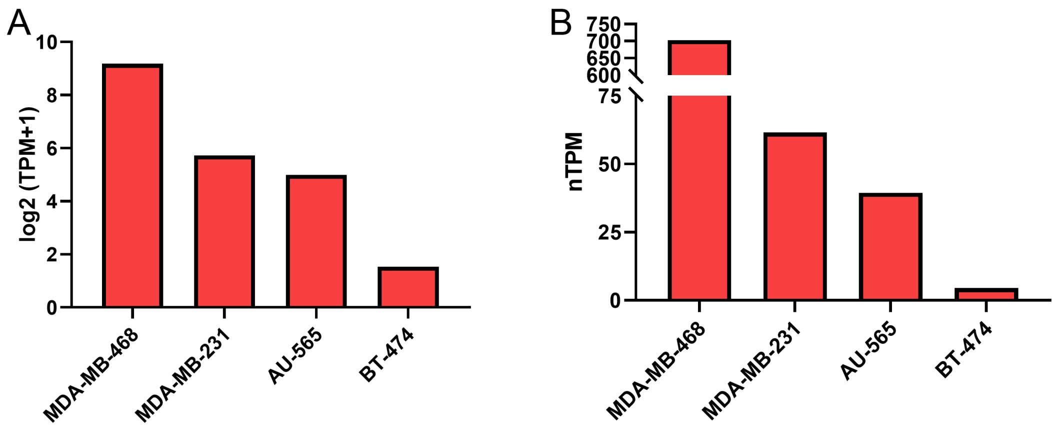


**Figure S4.** MDA-MB-468 cells express the highest levels of EGFR across the four cancer cell lines used in this study. EGFR gene expression data from the Depmap portal (A) or the Protein Atlas portal (B) for the cell lines as named, represented in transcripts per million (A: log TPM+1; B: normalised TPM).

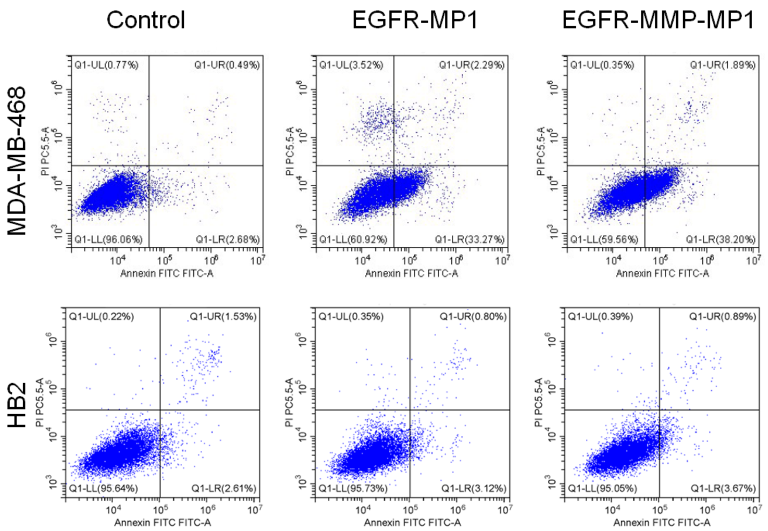


**Figure S5.** Representative primary data for Figure 3A. HB2 or MDA-MB-468 cells in 2D culture were treated with control, EGFR-MP1 (39.6μM) or EGFR-MMP-MP1 (56.6μM) for 24h. Apoptosis was quantified using Annexin V/PI staining and flow-cytometry (Annexin on the x-axis / PI on the y-axis). Cells in the two right quadrants were regarded as apoptotic.

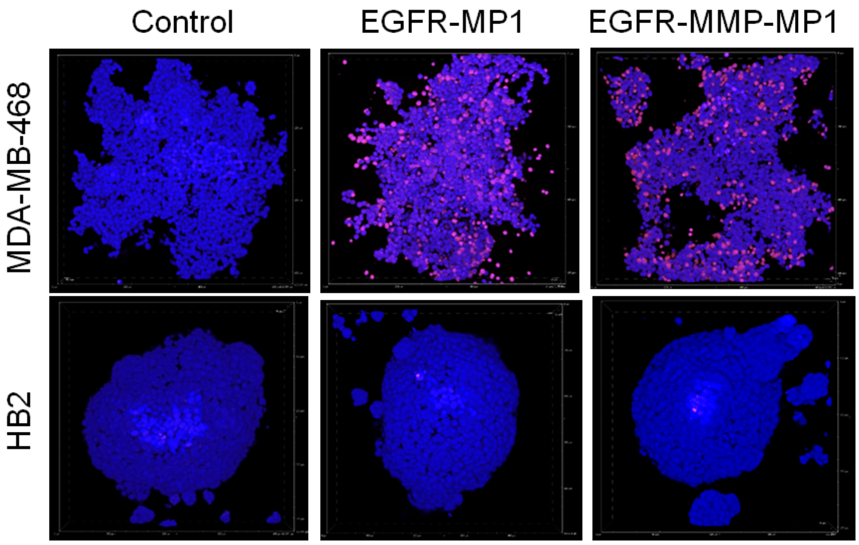


**Figure S6.** Representative primary data for Figure 3B. HB2 or MDA-MB-468 cells in 3D culture were treated with control, EGFR-MP1 (39.6μM) or EGFR-MMP-MP1 (56.6μM) for 24h. Cell survival was quantified by counting fluorescent cells after staining with PI/Hoechst 33342 by fluorescence microscopy. Red cells represented staining by PI (dead), whereas blue cells represented Hoechst 33342 staining allowing visualisation of the total cell population.

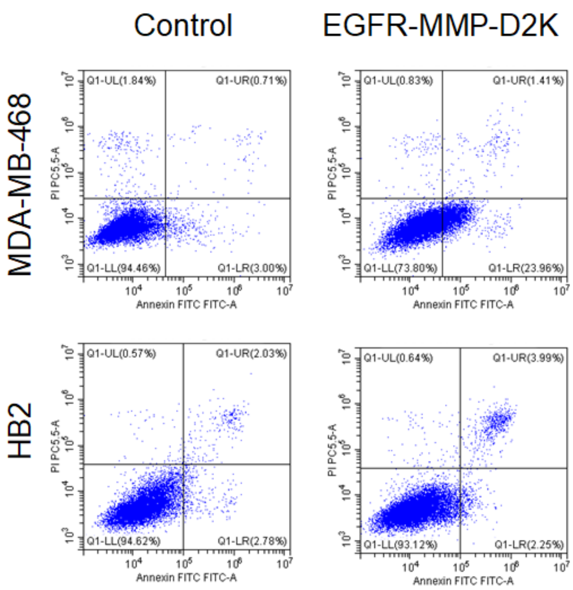


**Figure S7.** Representative primary data for Figure 4C. HB2 or MDA-MB-468 cells in 2D culture were treated with control or EGFR-MMP-D2K (32.1μM) for 24h. Apoptosis was quantified using Annexin V/PI staining and flow-cytometry (Annexin on the x-axis / PI on the y-axis). Cells in the two right quadrants were regarded as apoptotic.

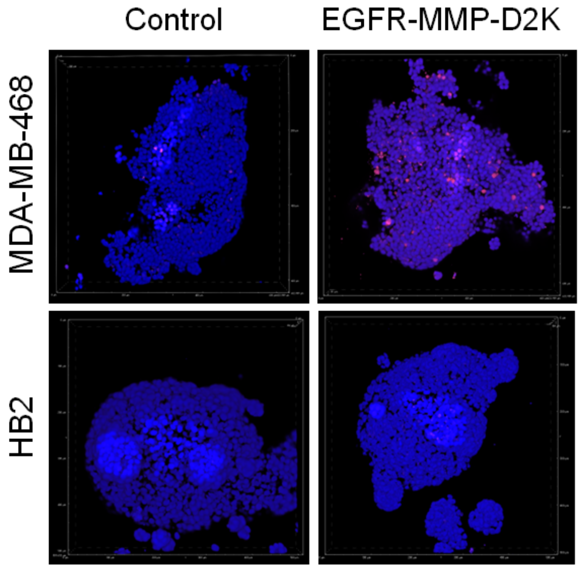


**Figure S8.** Representative primary data for Figure 4D. HB2 or MDA-MB-468 cells in 3D culture were treated with control or EGFR-MMP-D2K (32.1μM) for 24h. Cell survival was quantified by counting fluorescent cells after staining with PI/Hoechst 33342 by fluorescence microscopy. Red cells represented staining by PI (dead), whereas blue cells represented Hoechst 33342 staining allowing visualisation of the total cell population.
